# Admixture, Population Structure and *F*-statistics

**DOI:** 10.1101/028753

**Authors:** Benjamin M Peter

## Abstract

Many questions about human genetic history can be addressed by examining the patterns of shared genetic variation between sets of populations. A useful methodological framework for this purpose are *F*-statistics, that measure shared genetic drift between sets of two, three and four populations, and can be used to test simple and complex hypotheses about admixture between populations. Here, we put these statistics in context of phylogenetic and population genetic theory. We show how measures of genetic drift can be interpreted as branch lengths, paths through an admixture graph or in terms of the internal branches in coalescent trees. We show that the admixture tests can be interpreted as testing general properties of phylogenies, allowing us to generalize applications for arbitrary phylogenetic trees. Furthermore, we derive novel expressions for the *F*-statistics, which enables us to explore the behavior of *F*-statistic under population structure models. In particular, we show that population substructure may complicate inference.

**Author Summary:** For the analysis of genetic data from hundreds of populations, a commonly used technique are a set of simple statistics on data from two, three and four populations. These statistics are used to test hypotheses involving the history of populations, in particular whether data is consistent with the history of a set of populations forming a tree.

Here, we provide context to these statistics by deriving novel expressions and by relating them to approaches in comparative phylogenetics. These results are useful because they provide a straightforward interpretation of these statistics under many demographic processes and lead to simplified expressions. However, the result also reveals the limitations of *F*-statistics, in that population substructure may complicate inference.

## Introduction

For humans, whole-genome genotype data is now available for individuals from hundreds of populations [1,2], opening up the possibility to ask more detailed and complex questions about our history [3], and stimulating the development of new tools for the analysis of the joint history of these populations [4-9]. A simple and intuitive framework for this purpose that has quickly gained in popularity are the *F*-statistics, introduced by Reich *et al.* [4], and summarized in [5]. In that framework, inference is based on “shared genetic drift” between sets of populations, under the premise that shared drift implies a shared evolutionary history. Tools based on this framework have quickly become widely used in the study of human genetic history, both for ancient and modern DNA [1,10-13].

Some care is required with terminology, as the *F*-statistics *sensu* Reich *et al.* [4] are distinct, but closely related to Wright’s fixation indices [4,14], which are also often referred to as *F*-statistics. Furthermore, it is necessary to distinguish between statistics (quantities calculated from data) and the underlying parameters (which are part of the model, and typically what we want to estimate using statistics) [15].

In this paper, we will mostly discuss model parameters, and we will therefore refer to them as *drift indices.* The term *F*-*statistics* will be used when referring to the general framework introduced by Reich *et al.* [4], and Wright’s statistics will be referred to as *F_ST_* or *f*.

Most applications of the *F*-statistic-framework can be phrased in terms of the following six questions:

1. (Treeness test): Are populations related in a tree-like fashion? [4]
2. (Admixture test): Is a particular present day population descended from multiple ancestral populations? [4]
3. (Admixture ratio): What are the contributions from different population to a focal population [10].
4. (Number of founders): How many founder populations are there for a certain region? [1,11]
5. (Complex demography): How can mixtures and splits of population explain demography? [5,7]
6. (Closest relative): What is the closest relative to a contemporary or ancient population [16]

The demographic models under which these questions are addressed, and that motivated the drift indices, are called *population phylogenies* and *admixture graphs.* The population phylogeny (or population tree), is a model where populations are related in a tree-like fashion (Figure 1A), and it frequently serves as the null model for admixture tests. The branch lengths in the population phylogeny, correspond to genetic drift, so that a branch that is subtended by two different populations can be interpreted as the “shared” genetic drift between these populations. The alternative model is an admixture graph (Figure 1B), which extends the population phylogeny by allowing further edges that represent population mergers or a significant exchange of migrants.

**Figure 1.**
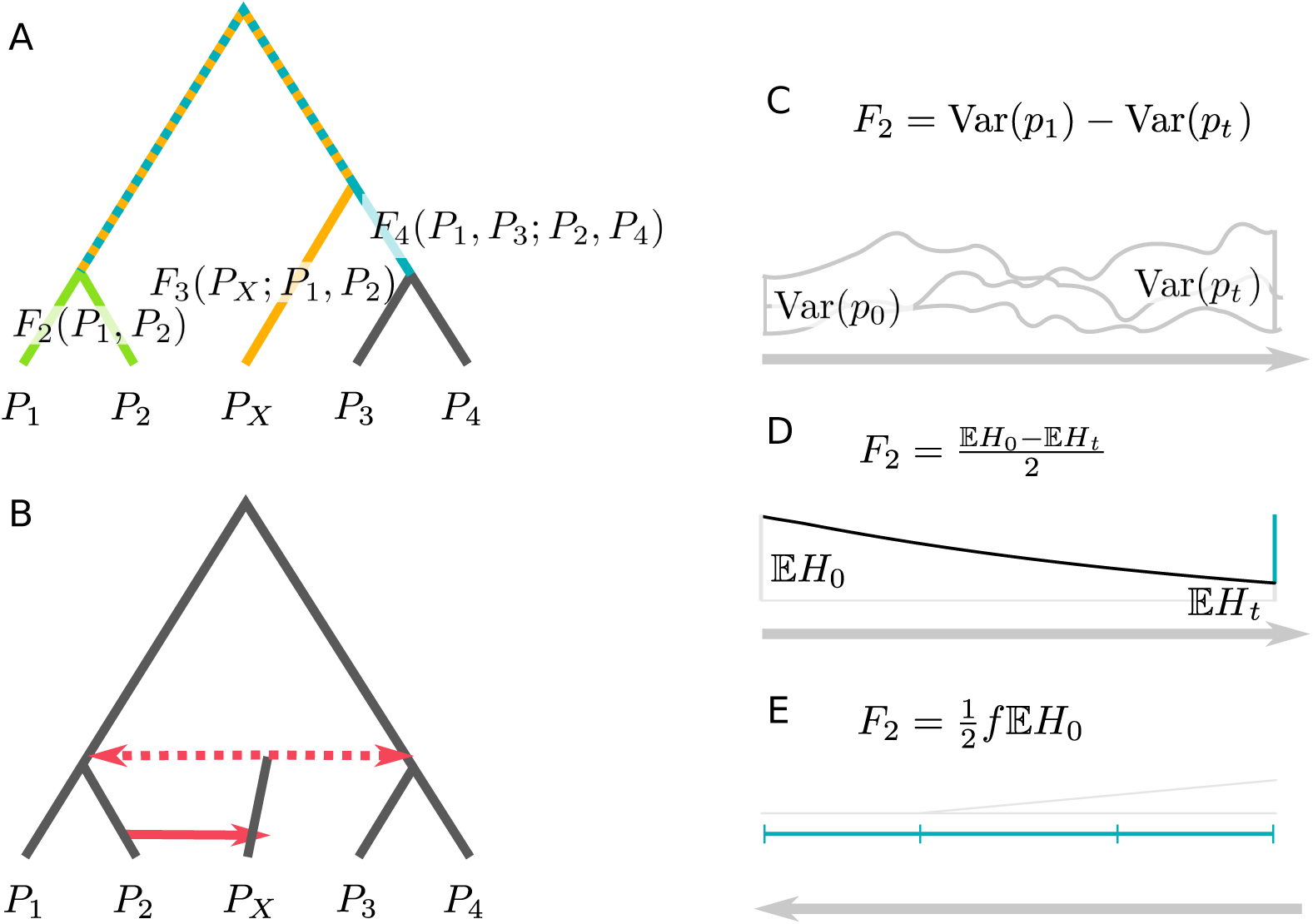
Schematics of gene genalogies, admixture graphs and measures of genetic drift. A: A population phylogeny with branches corresponding to *F*_2_ (green), *F*_3_ (yellow) and *F*_4_ (blue). The dotted branch is part of both *F*_3_ and *F*_4_. B: An Admixture graph, extends population phylogenies by allowing gene flow (red, full line) and admixture events (red, dotted). C-E: Interpretations of *F*_2_ in terms of allele frequency variances (C), heterozygosityies (D) and *f*, which can be interpreted as probability of coalescence of two lineages, or the probability that they are identical by descent.

The three *F*-statistics proposed by Reich *et al.* [4], labelled *F*_2_, *F*_3_ and *F*_4_, have simple interpretations under a population phylogeny: *F*_2_ corresponds to the path between two samples or vertices in the tree, whereas *F*_3_ and *F*_4_ can be interpreted as external and internal branches of the phylogeny, respectively (Figure 1A, [4]). In an admixture graph, there is no longer a single branch, and interpretations are more complex. However, *F*-statistics can be thought of as the proportion of genetic drift shared between populations [4].

The fundamental idea exploited in addressing all six questions outlined above is that under a tree model, branch lengths, and thus the drift indices, must satisfy some constraints [4,17,18]. The two most relevant constraints are that i) in a tree, all branches have positive lengths (tested using the *F*_3_-admixture test) and ii) in a tree with four leaves, there is at most one internal branch (tested using the *F*_4_-admixture test).

The goal of this paper is to give a broad overview on the theory, ideas and applications of *F*-statistics. Our starting point is a brief review on how genetic drift is quantified in general, and how it is measured using *F*_2_. We then propose an alternative definition of *F*_2_ that allows us to simplify some applications of *F*-statistics, and study them under a wide range of population structure models. We then review some basic properties of distance-based phylogentic trees, show how the admixture tests are interpreted in this context and evaluate their behavior. Many of the results we highlight here are implicit in classical [14,19-25] and more recent work [5-7], but often not explicitly stated, or given in a different context.

## Results & Discussion

In the next sections we will discuss the *F*-statistics, develop different interpretations and derive some useful expressions. Longer derivations are deferred to the Methods section. A graphical summary of the three interpretations of the statistics is given in Figure 2, and the main formulas are given in Table 1.

**Figure 2.**
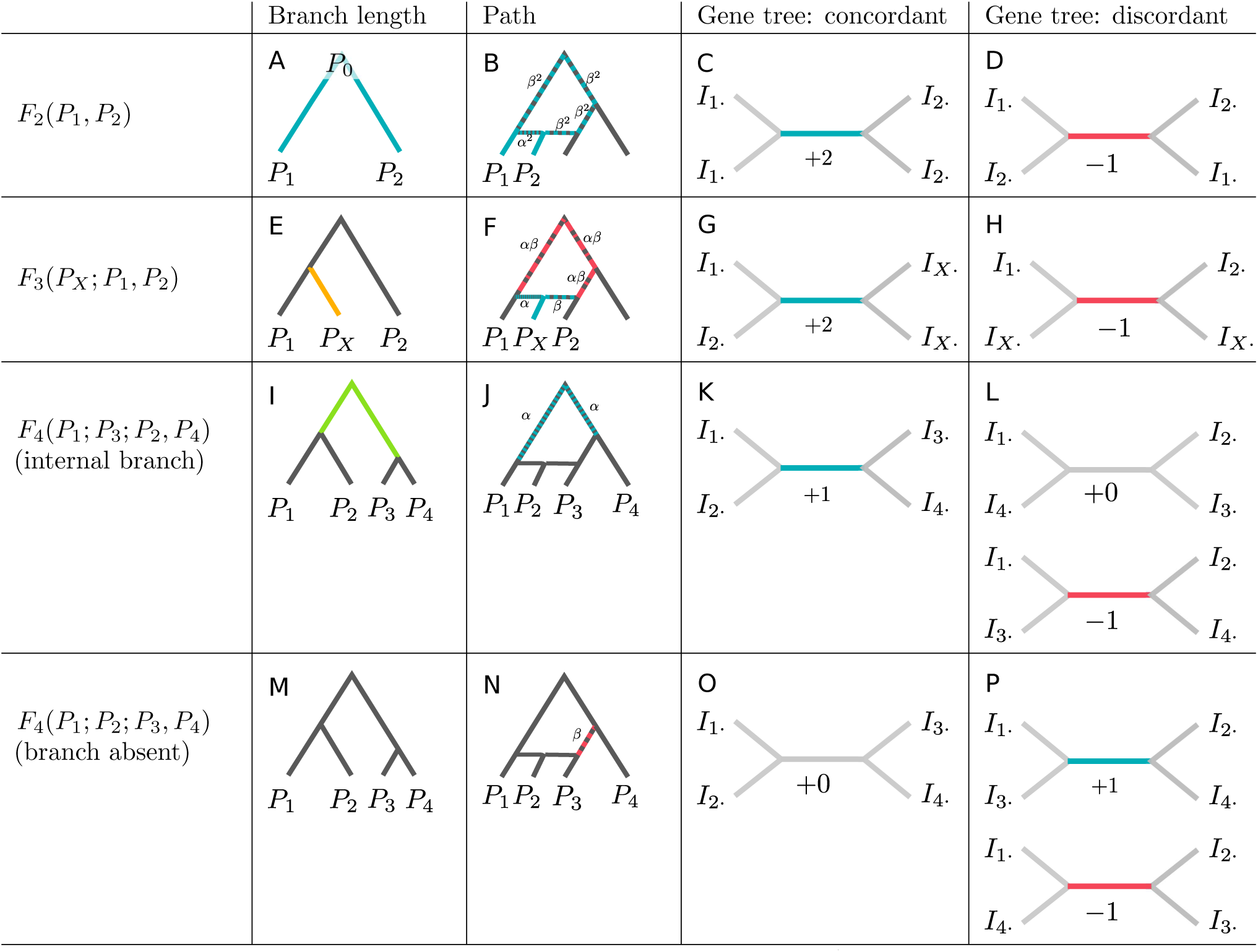
Interpretation of *F*-statistics. We can interpret the *F*-statistics i) as branch lengths in a population phylogeny (Panels A,E,I,M), the overlap of paths in an admixture graph (Panels B,F,J,N, see also Figure S1), and in terms of the internal branches of gene-genealogies (see Figures 3, S2 and S3). For gene trees consistent with the population tree, the internal branch contributes positively (Panels C,G,K), and for discordant branches, internal branches contribute negatively (Panels D,H) or zero (Panel L). For the admixture test, the two possible gene trees contribute to the statistic with different sign, highlighting the similarity to the *D*-statistic [10] and its expectation of zero in a symmetric model.

**Table 1.**
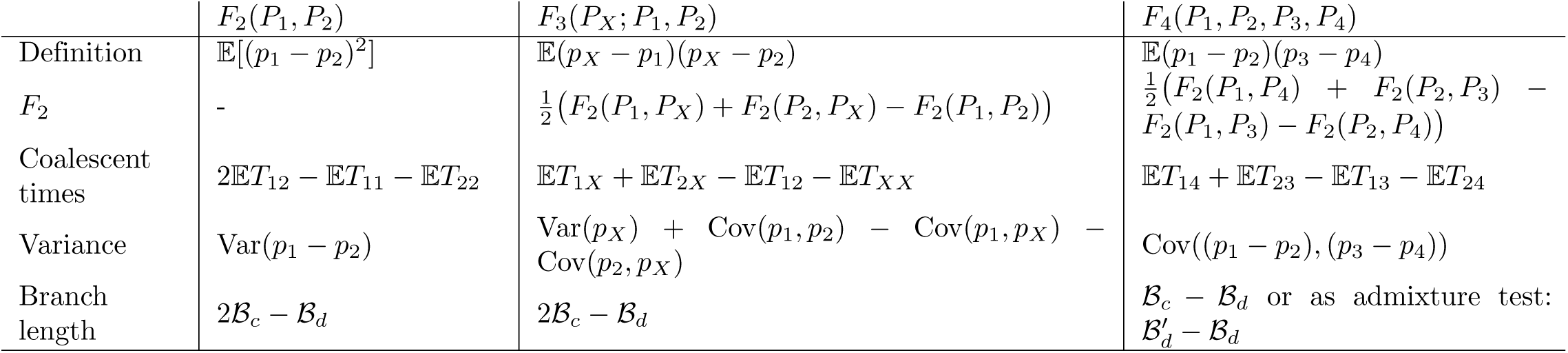
*F***-**statistics in terms of *F*_2_ or tree metrics, coalescent times and allele frequency variances. A constant of proportionality is omitted for coalescence times and branch lengths. Derivations for *F*_2_ are given in the main text, *F*_3_ and *F*_4_ are a simple result of combining Equations 20, 5 with 10b and 14b

| | $F_2(P_1, P_2)$ | $F_3(P_X; P_1, P_2)$ | $F_4(P_1, P_2, P_3, P_4)$ |
| --- | --- | --- | --- |
| Definition | $\mathbb{E}[(p_1 - p_2)^2]$ | $\mathbb{E}(p_X - p_1)(p_X - p_2)$ | $\mathbb{E}(p_1 - p_2)(p_3 - p_4)$ |
| $F_2$ | - | $\frac{1}{2}(F_2(P_1, P_X) + F_2(P_2, P_X) - F_2(P_1, P_2))$ | $\frac{1}{2}(F_2(P_1, P_4) + F_2(P_2, P_3) - F_2(P_1, P_3) - F_2(P_2, P_4))$ |
| Coalescent times | $2\mathbb{E}T_{12} - \mathbb{E}T_{11} - \mathbb{E}T_{22}$ | $\mathbb{E}T_{1X} + \mathbb{E}T_{2X} - \mathbb{E}T_{12} - \mathbb{E}T_{XX}$ | $\mathbb{E}T_{14} + \mathbb{E}T_{23} - \mathbb{E}T_{13} - \mathbb{E}T_{24}$ |
| Variance | $\text{Var}(p_1 - p_2)$ | $\text{Var}(p_X) + \text{Cov}(p_1, p_2) - \text{Cov}(p_1, p_X) - \text{Cov}(p_2, p_X)$ | $\text{Cov}((p_1 - p_2), (p_3 - p_4))$ |
| Branch length | $2\mathcal{B}_c - \mathcal{B}_d$ | $2\mathcal{B}_c - \mathcal{B}_d$ | $\mathcal{B}_c - \mathcal{B}_d$ or as admixture test: $\mathcal{B}'_d - \mathcal{B}_d$ |

Throughout this paper, we label populations as *P*_1_, *P*_2_,… *P_i_* Often, we will denote a potentially admixed population with *P_X_*, and an ancestral population with *P*_0_. The allele frequency *p_i_* is defined as the proportion of individuals in *P_i_* that carry a particular allele at a biallelic locus, and throughout this paper we will assume that all individuals are haploid. However, all results hold if instead of haploid individuals, we use a random allele of a diploid individual. If necessary, *t_i_* denotes the time when population *P_i_* is sampled. We focus on genetic drift only, and ignore the effects of mutation, selection and other evolutionary forces.

### Measuring genetic drift – *F*_2_

The first *F*-statistic we introduce is *F*_2_, whose purpose is simply to measure genetic dissimilarity or how much genetic drift occurred between two populations. For populations *P*_1_ and *P*_2_, *F*_2_ is defined as [4]

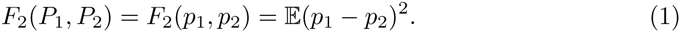

The expectation is with respect to the evolutionary process, but in practice *F*_2_ is estimated from hundreds of thousands of loci across the genome [5], which are assumed to be non-independent replicates of the evolutionary history of the populations.

Why is *F*_2_ a useful measure of genetic drift? As it is generally infeasible to observe the changes in allele frequency directly, we assess the effect of drift indirectly, through its impact on genetic diversity. In general, genetic drift is quantified in terms of i) the variance in allele frequency, ii) heterozygosity, iii) probability of identity by descent iv) correlation (or covariance) between individuals and v) the probability of coalescence (two lineages having a common ancestor).

#### Single population

To make these measures of drift explicit, we assume a single population, measured at two time points (*t*_0_ ≤ *t_t_*), and label the two samples *P*_0_ and *P_t_*. Then *F*_2_(*P*_0_, *P_t_*) can be interpreted in terms of the variance of allele frequencies (Figure 1C)

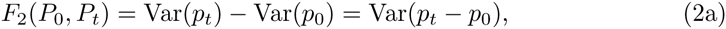

the expected decrease in heterozygosity *H_t_*, between the two sample times (Figure 1D):

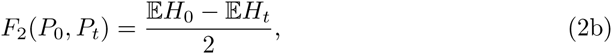

and in terms of the inbreeding coefficient *f*, which can be interpreted as the probability of two individuals in *P_t_* descend from the same ancestor in *P*_0_, or, equivalently, the probability that two samples from *P_t_* coalesce before *t*_0_. (Figure 1E, [26]):

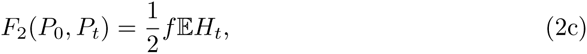

Rearranging Equation 2b, we find that 2*F*_2_ simply measures the absolute decrease of heterozyosity through time

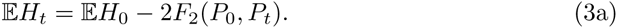

If we assume that we know *p*_0_ and therefore Var(*p*_0_) is zero, we can combine 2a and 2b and obtain

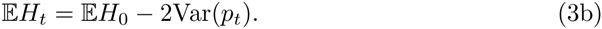

Similarly, using equations 2b and 2c we obtain an expression in terms of *f*.

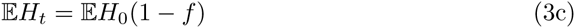

#### Pairs of populations

Equations 3b and 3c describing the decay of heterozygosity are – of course – well known by population geneticists, having been established by Wright [14]. In structured populations, very similar relationships exist when we compare the number of heterozygotes expected from the overall allele frequency, *H_obs_* with the number of heterozygotes present due to differences in allele frequencies between populations *H_exp_* [14,19].

In fact, already Wahlund showed that for a population made up of two subpopulations with equal proportions, the proportion of heterozygotes is reduced by

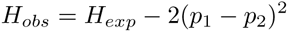

from which it is easy to see that

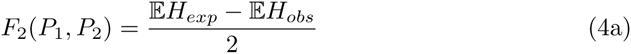

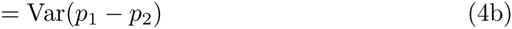

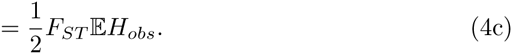

This last equation served as the original motivation of *F*_2_ [4], which was first defined as a numerator of *F_ST_*.

#### Justification for *F*_2_

Our preceding arguments show how the usage of *F*_2_ for both single and structured populations can be justified by the similar effects on heterozygosity and allele frequency variance *F*_2_ measures. However, what is the benefit of using *F*_2_ instead of the established inbreeding coefficient *f* and fixation index *F_ST_*? A conceptual way to approach this is by recalling that Wright motivated *f* and *F_st_* as *correlation coefficients* between alleles [14,27]. This has the advantage that they are easy to interpret, as, e.g. *F_ST_* = 0 implies panmixia and *F_ST_* = 1 implies complete divergence between subpopulations. In contrast, *F*_2_ depends on allele frequencies and is highest for intermediate frequency alleles. However, *F*_2_ has an interpretation as a *covariance*, making it simpler and mathematically more convenient to work with. In particular, variances and covariances are frequently partitioned into components due to different effects using techniques such as analysis of variance and analysis of covariance (e.g. [25]).

#### *F*_2_ as branch length

Reich et al. [4,5] proposed to partition “drift” (as we established, characterized by allele frequency variance, or decrease in heterozygosity) between different populations into contribution on the different branches of a population phylogeny. This model has been studied by Cavalli-Sforza & Edwards [20] and Felsenstein [21] in the context of a Brownian motion process. In this model, drift on independent branches is assumed to be independent, meaning that the variances can simply be added. This is what is referred to as the *additivity property* of *F*_2_ [5].

To illustrate the additivity property, consider two populations *P*_1_ and *P*_2_ that split recently from a common ancestral population *P*_0_ (Figure 2A). In this case, *p*_1_ and *p*_2_ are independent conditional on *p*_0_, and therefore Cov(*p*_1_, *p*_2_) = Var(*p*_0_). Then, using 2a and 4b,

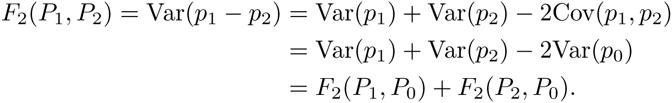

Alternative proofs of this statement and more detailed reasoning behind the additivity assumption can be found in [4, 5, 20, 21].

For an admixture graph, we cannot use this approach as lineages are not independent. Reich *et al.* [4] approached this by conditioning on the possible population trees that are consistent with an admixture scenario. In particular, they proposed a framework of counting the possible *paths* through the graph [4,5]. An example of this representation for *F*_2_ in a simple admixture graph is given in Figure S1, with the result summarized in Figure 2B. Detailed motivation behind this visualization approach is given in Appendix 2 of [5]. In brief, the reasoning is as follows: We write 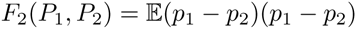, and interpret the two terms in parentheses as two paths between *P*_1_ and *P*_2_, and *F*_2_ as the overlap of these two paths. In a population phylogeny, there is only one possible path, and the two paths are always the same, therefore *F*_2_ is the sum of the length of all the branches connecting the two populations. However, if there is admixture, as in Figure 2B, both paths choose independently which admixture edge they follow. With probability α they will go left, and with probability *β* = 1 − *α* they go right. Thus, *F*_2_ can be interpreted by enumerating all possible choices for the two paths, resulting in three possible combinations of paths on the trees (Figure S1), and the branches included will differ depending on which path is chosen, so that the final *F*_2_ is made up average of the path overlap in the topologies, weighted by the probabilities of the topologies.

However, one drawback of this approach is that it scales quadratically with the number of admixture events, making calculations cumbersome when the number of admixture events is large. More importantly, this approach is restricted to panmictic subpopulations, and cannot be used when the population model cannot be represented as a weighted average of trees.

#### Gene tree interpretation

For this reason, we propose to redefine *F*_2_ using coalescence theory [28]. Instead of allele frequencies on a fixed admixture graph, we track the ancestors of a sample of individuals, tracing their history back to their most recent common ancestor. The resulting tree is called a *gene tree* (or coalescent tree). Gene trees vary between loci, and will often have a different topology from the population phylogeny, but they are nevertheless highly informative about a population’s history. Moreover, expected coalescence times and expected branch lengths are easily calculated under a wide array of neutral demographic models.

In a seminal paper, Slatkin [24] showed how *F_ST_* can be interpreted in terms of the expected coalescence times of gene trees:

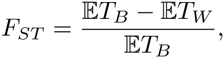

where 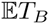 and 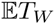 are the expected coalescence times of two lineages sampled in two different and the same population, respectively.

Unsurprisingly, given the close relationship between *F*_2_ and *F_ST_*, we may obtain a similar expression for *F*_2_(*P*_1_, *P*_2_):

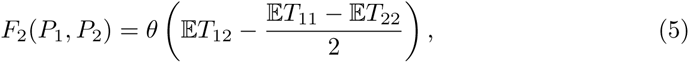

where *θ* is a scaled mutation parameter, *T*_12_ is the expected coalescence time for one lineage each sampled from populations *P*_1_ and *P*_2_, and *T*_11_, *T*_22_ are the expected coalescence times for two samples from the *P*_1_ and *P*_2_, respectively. Unlike *F_ST_*, the mutation parameter *θ* does not cancel. However, for most applications, the absolute magnitude of *F*_2_ is of little interest, as we are only interested if a sum of *F*_2_-values is significantly different from zero, significantly negative, or we are comparing *F*-statistics with the same *θ* [4]. For this purpose, we may regard *θ* as a constant of proportionality and largely ignore its effect.

For estimation, the average number of pairwise differences *π_ij_* is a commonly used estimator for *θT_ij_* [29]. Thus, we can write the estimator for *F*_2_ as

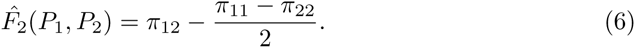

This estimator of *F*_2_ is numerically equivalent to the unbiased estimator proposed by [4] in terms of the sample allele frequency 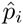 and the sample size *n_i_* (Equation 10 in the Appendix of [4]):

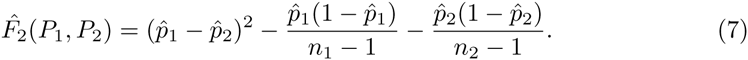

However, the modelling assumptions are different: The original definition only considered loci that were segregating in the the ancestral population; loci not segregating there were discarded. Since ancestral populations are usually unsampled, this is often replaced by ascertainment in an outgroup [5,7]. In contrast, Equation 6 assumes that all markers are used, which is more convenient for sequence data.

#### Gene tree branch lengths

An important feature of Equation 5 is that it only depends on the coalescence times between pairs of lineages. Thus, we may fully characterize *F*_2_ by considering a sample of size four, with two random individuals taken from each population, as this allows us to study the joint distribution of *T*_12_, *T*_11_ and *T*_22_. For a sample of size four with two pairs, there are only two possible unrooted tree topologies. One, where the lineages from the same population are more closely related to each other (called *concordant* topology, 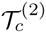) and one where lineages from different populations coalesce first (which we will refer to as *discordanat* topology 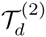). The superscripts refers to the topologies being for *F*_2_, and we will discard them in cases where no ambiguity arises.

Thus, we can condition on the topology, and ask how *F*_2_ depends on the topology:

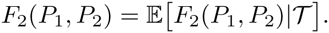

One way to do that is for each topology, consider each of the pairwise differences in Equation 5 separately, and then add the branches (see Figure 3 for a graphical representation).

**Figure 3.**
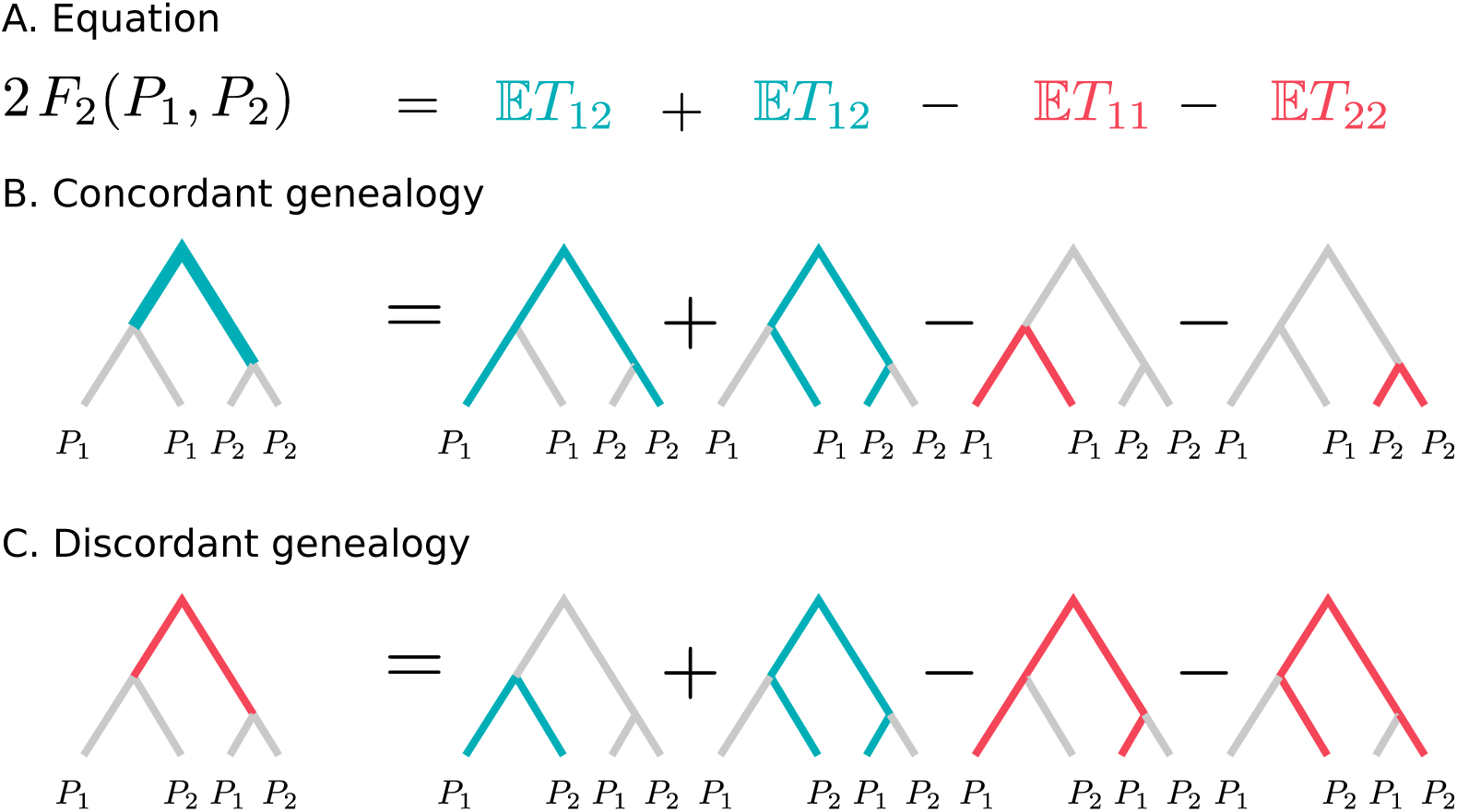
Schematic explanation how *F*_2_ behaves conditioned on gene tree. Blue terms and branches correspond to positive contributions, whereas red branches and terms are subtracted. Labels represent individuals randomly sampled from that population. We see that external branches cancel out, so only the internal branches have non-zero contribution to *F*_2_. In the concordant genealogy (Panel B), the contribution is positive (with weight 2), and in the discordant genealogy (Panel C), it is negative (with weight 1). The mutation rate as constant of proportionality is omitted.

We see that in both topologies, only the internal branch has a non-zero impact on *F*_2_, and the contribution of the external branches cancels out. The external branch leading to a sample from *P*_1_, for example, is included with 50% probability in *T*_12_, but will always be included in *T*_11_, so these two terms negate the effect of that branch. The internal branch of 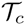 will contribute with a factor of *a_c_* = 2 to *F*_2_, since the internal branch is added twice in Figure 3B. In contrast, the length of the internal branch of 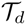 is subtracted from *F*_2_, with coefficient *a_d_* = −1. Thinking of *F*_2_ as a distance between population that is supposed to be large when the populations are very different from each other, this makes intuitive sense: if the populations are closely related we expect to see 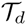 relatively frequently, and *F*_2_ will be low. However, if the populations are more distantly related, then 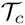 will be most common, and *F*_2_ will be large.

An interesting way to represent *F*_2_ is therefore in terms of the internal branches over all possible gene genealogies. Let us denote the unconditional average length of the internal branch of *T_c_* as 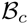. Similarly, we denote the average length of the internal branch in 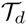 as 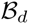.

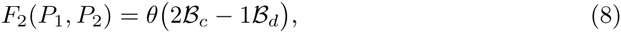

with the coefficients *a_c_* = 2, *a_d_* = 1. A graphical summary of this is given in Figure 2C-D. As a brief sanity check, we can consider the case of a population without structure. In this case, the branch length is independent of the topology and 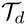 is twice as likely as 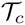. In this case, we see immediately that *F*_2_ will be zero, as expected when there is no difference between topologies.

## Testing treeness

In practical cases, we often have dozens or even hundreds of populations [2,5,12], and we want to infer where and between which populations admixture occurred. Using *F*-statistics, the approach is to interpret *F*_2_(P_1_, *P*_2_) as a measure of dissimilarity between *P*_1_ and *P*_2_, as a large *F*_2_-value implies that populations are highly diverged. Thus, we calculate all pairwise *F*_2_ indices between populations, combine them into a *dissimilarity matrix,* and ask if that matrix is consistent with a tree.

One way to approach this question is by using phylogenetic theory: Many classical algorithms have been proposed that use a measure of dissimilarity to generate a tree [18,30-32], and what properties a general dissimilarity matrix needs to have in order to be consistent with a tree [17,22], in which case the matrix is also called a *tree metric* [18].

There are two central properties for a dissimilarity matrix to be consistent with a tree: The first property is that all edges in a tree have positive length. This is strictly not necessary for phylogenetic trees, and some algorithms may return negative branch lengths [31]; however, since in our case branches have an interpretation of genetic drift, it is clear that negative genetic drift is biologically meaningless, and therefore negative branches should be interpreted as a violation of the modelling assumptions and hence treeness.

The second property of a tree metric that we require is a bit more involved: A dissimilarity matrix (written in terms of F_2_) is consistent with a tree if for any four populations *P_i_*, *P_j_*, *P_k_* and *P_l_*,

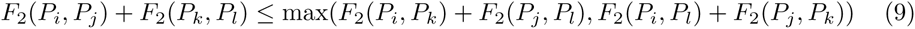

that is, if we compare the sums of all possible pairs of distances between populations, then two of these sums will be the same, and no smaller than the third. This theorem, due to Buneman [17,33] is called the four-point condition or sometimes, more modestly, the “fundamental theorem of phylogenetics”. A proof can be found in Chapter 7 of [18].

In terms of a tree, this statement can be understood by noticing that on a tree, two of the pairs of distances will include the internal branch, whereas the third one will not, and therefore be shorter, or the same length for a topology with no internal branch. Thus, the four-point condition can be informally rephrased as “for any four taxa, a tree has at most one internal branch”.

Why are these properties useful? It turns out that the admixture tests based on *F*-statistics can be interpreted as tests of these properties: The *F*_3_-test can be interpreted as a test for the positivity of a branch; and the *F*_4_ as a test of the four-point condition. Thus, we can interpret the working of the two test statistics in terms of fundamental properties of phylogenetic trees, with the immediate consequence that they can be applied as treeness-tests for arbitrary dissimilarity matrices.

An early test of treeness, based on a likelihood ratio, was proposed by Cavalli-Sforza & Piazza [22]: They compare the likelihood of the observed *F*_2_-matrix matrix to that induced by the best fitting tree (assuming Brownian motion), rejecting the null hypothesis if the tree-likelihood is much lower than that of the empirical matrix. In practice, however, finding the best-fitting tree is a challenging problem, especially for large trees [32] and so the likelihood test proved to be difficult to apply. From that perspective, the *F*_3_ and *F*_4_-tests provide a convenient alternative: Since treeness implies that all subsets of taxa are also trees, the ingenious idea of Reich *et al.* [4] was that rejection of treeness for subtrees of size three (for *F*_3_) and four (for *F*_4_) is sufficient to reject treeness for the entire tree [4]. Furthermore, tests on these subsets also pinpoint the populations involved in the non-tree-like history.

### *F*_3_

In the previous section, we showed how *F*_2_ can be interpreted as a branch length, an overlap of paths or in terms of gene trees (Figure 2). Furthermore, we derived expressions in terms of coalescent times, allele frequency variances and internal branch lengths of gene trees. We now derive analogous results for *F*_3_.

Reich *et al.* [4] defined *F*_3_ as:

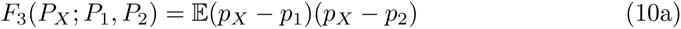

with the goal to test whether *P_X_* is admixed. Recalling the path interpretation detailed in [5], *F*_3_ can be interpreted as the shared portion of the paths from *P_X_* to *P*_1_ with the path from *P_X_* to *P*_1_. In a population phylogeny (Figure 2E) this corresponds to the branch between *P_X_* and the internal node. Equivalently, *F*_3_ can also be written in terms of *F*_2_ [4]:

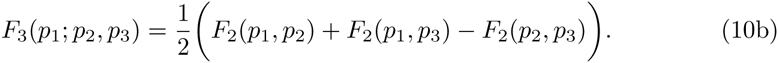

If we replace *F*_2_ in Equation 10b with an arbitrary tree metric, Equation 10b is known as the Gromov product [18] in phylogenetics. The Gromov product is a commonly used operation in classical phylogenetic algorithms to calculate the length of the portion of a branch shared bewtween *P*_1_ and *P*_2_ [21,30,31]: consistent with the notion that *F*_3_ is the length of an external branch in a phylogeny.

In an admixture graph, there is no longer a single external branch; instead we again have to consider all possible trees, and *F*_3_ is the (weighted) average of paths through the admixture graph (Figure 2F).

Combining Equations 5 and 10b, we find that *F*_3_ can be written in terms of expected coalescence times as

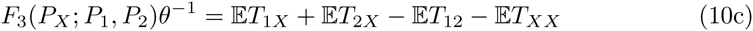

Similarly, we may obtain an expression for the variance by combining Equation 20 with 10b, and find that

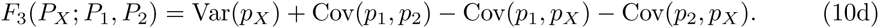

This result can also be found in [6].

#### Outgroup-*F*_3_ statistics

A simple application of the interpretation of *F*_3_ as a shared branch length are the “outgroup”-*F*_3_-statistics proposed by [16]. For an unknown population *P_U_*, they wanted to find the most closely related population from a panel of *k* extant populations {*P_i_*, *i* = 1, 2,… *k*}. They did this by calculating *F*_3_(*P_O_*, *P_U_*, *P_i_*), where *P_O_* is an outgroup population that was assumed widely diverged from *P_U_* and all populations in the panel. This measures the shared drift (or shared branch) of *P_U_* with the populations from the panel, and high *F*_3_-values imply close relatedness.

However, using Equation 10c, we see that the outgroup-*F*_3_-statistic is

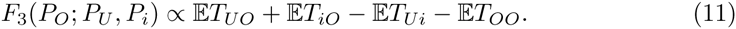

Out of these four terms, 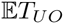 and 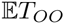 do not depend on the panel. Furthermore, if *P_O_* is truly an outgroup, then all 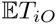 should be the same, as pairs of individuals from the panel population and the outgroup can only coalesce once they are in the joint ancestral population. Therefore, only the term 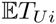 is expected to vary between different panel populations, suggesting that using the number of pairwise differences, *π_Ui_*, is largely equivalent to using *F*_3_ (*P_O_*; *P_U_*, *P_i_*). We confirm this in Figure 4A, where we calculate outgroup-*F*_3_ and *π_Ui_* for a set of increasingly divergent populations. Linear regression confirms the visual picture that *π_Ui_* has a higher correlation with divergence time (*R*^2^ = 0.75) than *F*_3_ (*R*^2^ = 0.49).

**Figure 4.**
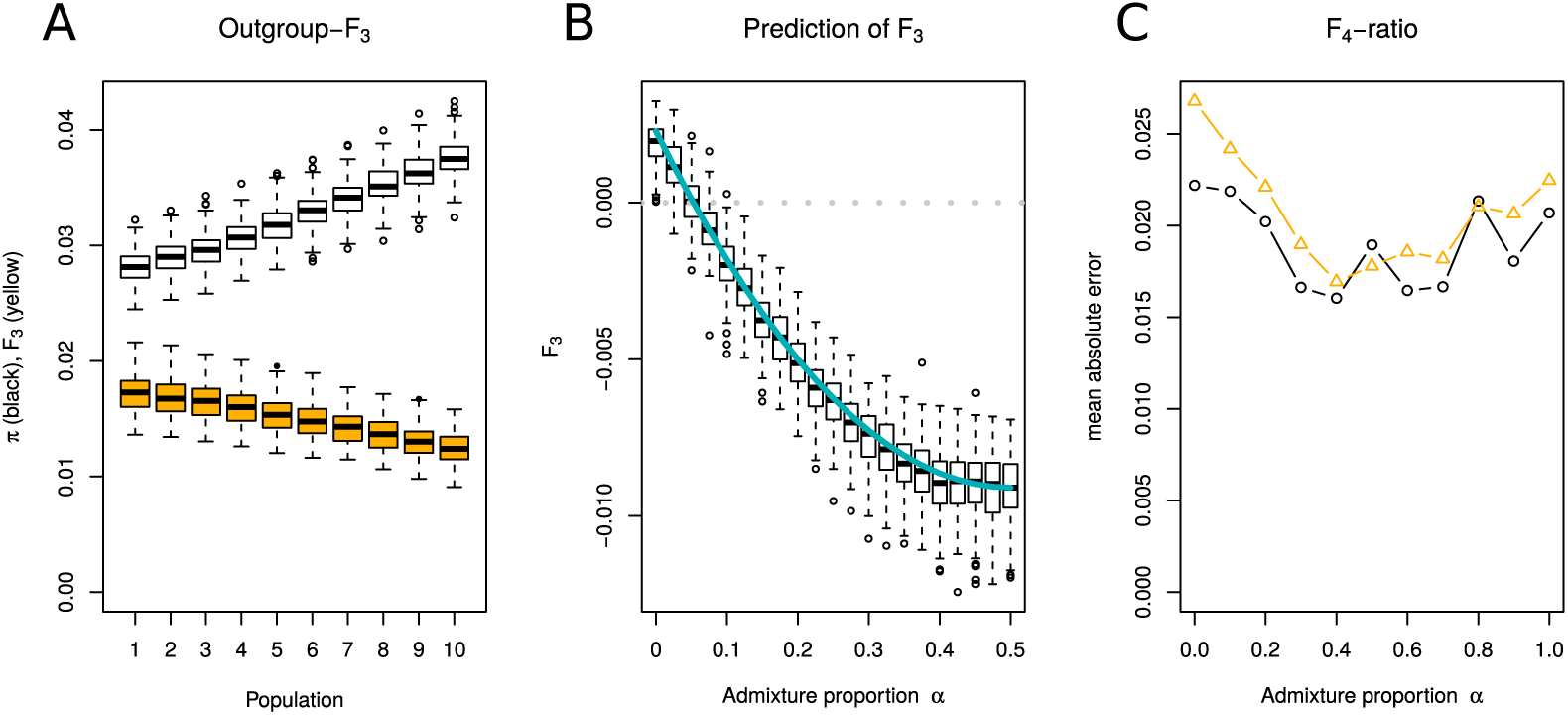
Simulation results. A: Outgroup-*F*_3_ statistics (yellow) and *π_iU_* (white) for a panel of populations with linearly increasing divergence time. B: Simulated (boxplots) and predicted(blue) *F*_3_-statistics under a simple admixture model (main text). C: Comparison of *F*_4_-ratio (yellow, Equation 17) and ratio of differences (Equation 19 black)

#### *F*_3_ admixture test

However, the main motivation of defining *F*_3_ has been as an admixture test [4]. In this context, the null hypothesis is that *F*_3_ is non-negative, i.e. we are testing if the data is consistent with a phylogenetic tree that has positive edge lengths. If this is not the case, we reject the tree model for the more complex admixture graph. From Figure 2F, we see that drift on the path on the internal branches (red) contribute negatively to *F*_3_. If these branches are long enough compared to the branch after the admixture event (blue), then *F*_3_ will be negative. For the simplest scenario where *P_X_* is admixed bewteen *P*_1_ and *P*_2_, Reich et al. [4] provided a condition when this is the case (Equation 20 in Supplement 2 of [4]). However, since this condition involves *F*-statistics with internal, unobserved populations, it is not easily applicable. We can obtain a more useful condition using gene trees:

In the simplest admixture model, an ancestral population splits into *P*_1_ and *P*_2_ and time *t_r_*. At time *t*_1_, the populations mix to form *P_X_*, such that with probability *α*, individuals in *P_X_* descend from individuals from *P*_1_, and with probability (1 − *α*), they descend from *P*_2_. In this case, *F*_3_(*P_X_*; *P*_1_, *P*_2_) is negative if

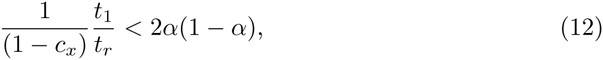

where *c_x_* is the probability two individuals sampled in *P_X_* have a common ancestor before *t*_1_. For a constant sized population of size 1, 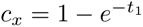. We see that power of *F*_3_ to detect admixture increases the closer they get to fifty percent, and that it only depends on the ratio between the original split and the secondary contact, and coalescence events that happen in *P_X_*.

We obtain a more general condition for negativity of *F*_3_ by considering the internal branches of the possible gene tree topologies, as we did for *F*_2_. Note that Equation 10c includes 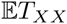, implying that we need two individuals from *P_X_*, but only one each from *P*_1_ and *P*_2_ to study the joint distribution of all terms in (10c). The minimal case is therefore contains again just four samples (Figure S2).

Furthermore, *P*_1_ and *P*_2_ are exchangeable, and thus we can again consider just two unrooted genealogies, a concordant one 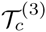 where the two lineages from *P_X_* are most closely related, and a discordant genealogy 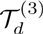 where the lineages from *P_X_* merge first with the other two lineages. A similar argument as that for *F*_2_ shows (presented in Figure S2) that *F*_3_ can be written as a function of just the internal branches in the topologies:

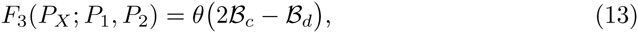

where 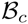 and 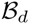 are the lengths of the internal branches in 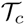 and 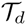, respectively, and similar to *F*_2_, they have coefficients *a_c_* = 2 and *a_d_* = −1. Again, if we do the sanity check of all samples coming from a single, randomly mating population, then 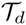 is again twice as likely as 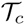, and all branches are expected to have the same length. Thus *F*_3_ is zero, as expected. However, for *F*_3_ to be negative, we see that 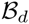 needs to be more than two times longer than 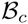. Thus, *F*_3_ can be seen as a test whether mutations that agree with the population tree are more common than mutations that disagree with it.

We performed a small simulation study to test the accuracy of Equation 12. Parameters were chosen such that *F*_3_ has a negative expectation for *α* > 0.05 (grey dotted line in Figure 4B), so simulations on the left of that line have positive expectation, and samples on the right are true positives. We find that our predicted *F*_3_ fits very well with the simulations (Figure 4B).

### *F*_4_

The second admixture statistic, *F*_4_, is defined as [4]

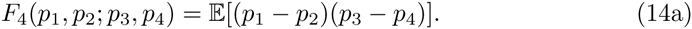

Similarly to *F*_3_, *F*_4_ can be written as a linear combination of *F*_2_:

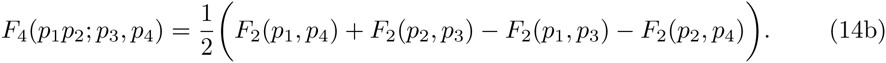

Equations giving *F*_4_ in terms of pairwise coalescence times and as a covariance are given in Table 1.

As four populations are involved, there are 4! = 24 possible ways of arranging the arguments in Equation 14a. However, there are four possible permutations of arguments that will lead to identical values, leaving only six unique *F*_4_-values for any four populations. Furthermore, these six values come in pairs that have the same absolute value, and a different sign, leaving only three unique absolute values, which correspond to the tree possible tree topologies. Thus, we may always find a way of writing *F*_4_ such that the statistic is non-negative (i. e. *F*_4_(*P*_1_, *P*_2_; *P*_3_, *P*_4_) = −*F*_4_(*P*_1_, *P*_2_; *P*_4_, *P*_3_)). Out of these three, one *F*_4_ can be written as the sum of the other two, leaving us with just two independent possibilities:

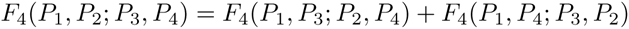

As we did for *F*_3_, we can generalize Equation 14b by replacing *F*_2_ with an arbitrary tree metric. In this case, Equation 14b is known as a tree split [17], as it measures the length of the overlap of the branch lengths between the two pairs (*P*_1_, *P*_2_) and (*P*_3_, *P*_4_). Tree splits have the property that if there exists a branch “splitting” the populations such that the first and third argument are on one side of the branch, and the second and fourth are on the other side (Figure 61), then it corresponds to the length of that branch. If no such branch exists, then *F*_4_ will be zero.

This can be summarized by the four-point condition [17,33], or, informally, by noting that any four populations will have at most one internal branch, and thus one of the three *F*_4_-values will be zero, and the other two will have the same value. Therefore, one *F*_4_-index has an interpretation as the internal branch in a genealogy, and the other can be used to test if the data corresponds to a tree. In Figure 2, the third row (Panels I-L) correspond to the internal branch, and the last row (Panels M-P) to the “zero”-branch.

Thus, in the context of testing for admixture, by testing that *F*_4_ is zero we check whether there is in fact only a single internal branch, and if that is not the case, we reject a population phylogeny for an admixture graph.

Evaluating *F*_4_ in terms of gene trees and their internal branches, we have to consider the three different possible gene tree topologies, and depending on if we want to estimate a branch length or do an admixture test, they are interpreted differently.

For the branch length, we see that the gene tree corresponding to the population tree has a positive contribution to *F*_4_, and the other two possible trees have a zero and negative contribution, respectively (Figure S3). Since the gene tree corresponding to the population tree is expected to be most frequent, *F*_4_ will be positive, and we can write

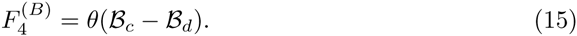

This equation is slightly different than those for *F*_2_ and *F*_3_, where the coefficient for the discordant genealogy was half that for the concordant genealogy. Note, however, that we have two discordant genealogies, and *F*_4_ only measures one of them. Under a tree, both discordant genealogies are equally likely [34], and thus the expectation of *F*_4_ will be the same.

In contrast, for the admixture test statistic, the contribution of the concordant genealogy will be zero, and the discordant genealogies will contribute with coefficients −1 and +1, respectively. Under the population phylogeny, these two gene trees will be equally likely [28], and thus the expectation of *F*_4_ as a test statistic

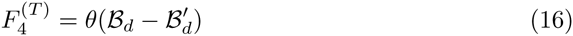

is zero under the null hypothesis. Furthermore, we see that the statistic is closely related to the ABBA-BABA or *D*-statistic also used to test for admixture [10,34], which includes a normalization term, and in our notation is defined as,

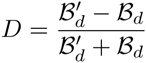

but otherwise tests the exact same hypothesis.

### *F*_4_-as a branch

#### Rank test

Two major applications of *F*_4_ use its interpretation as a branch length. First, we can use the rank of a matrix of all *F*_4_-statistics to obtain a lower bound on the number of admixture events required to explain data [11]. The principal idea of this approach is that the number of internal branches in a genealogy is bounded to be at most *n* − 3 in an unrooted tree. Since each *F*_4_ corresponds to a sum of internal branches, all *F*_4_-indices should be sums of *n* − 3 branches, or *n* – 3 independent components. This implies that the rank of the matrix (see e.g. Section 4 in [35]) is at most *n* – 3, if the data is consistent with a tree. However, admixture events may increase the rank of the matrix, as they add additional internal branches [11]. Therefore, if the rank of the matrix is *r*, the number of admixture events is at least *r* − *n* + 3.

One issue is that the full *F*_4_-matrix has size 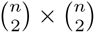, and may thus become rather large. Furthermore, in many cases we are only interested in admixture events in a certain part of the phylogeny. To estimate the number of admixture events on a particular branch of the phylogeny, Reich *et al.* [11], proposed to find two sets of test populations *S*_1_ and *S*_2_, and two reference populations for each set *R*_1_ and *R*_2_ that are presumed unadmixed (see Figure 5A). Assuming a phylogeny, all *F*_4_(*S*_1_, *R*_1_; *S*_2_, *R*_2_) will measure the length of the branch absent from Figure 5A, und should be zero, and the rank of the matrix of all *F*_4_ of that form reveals the number of branches of that form.

**Figure 5.**
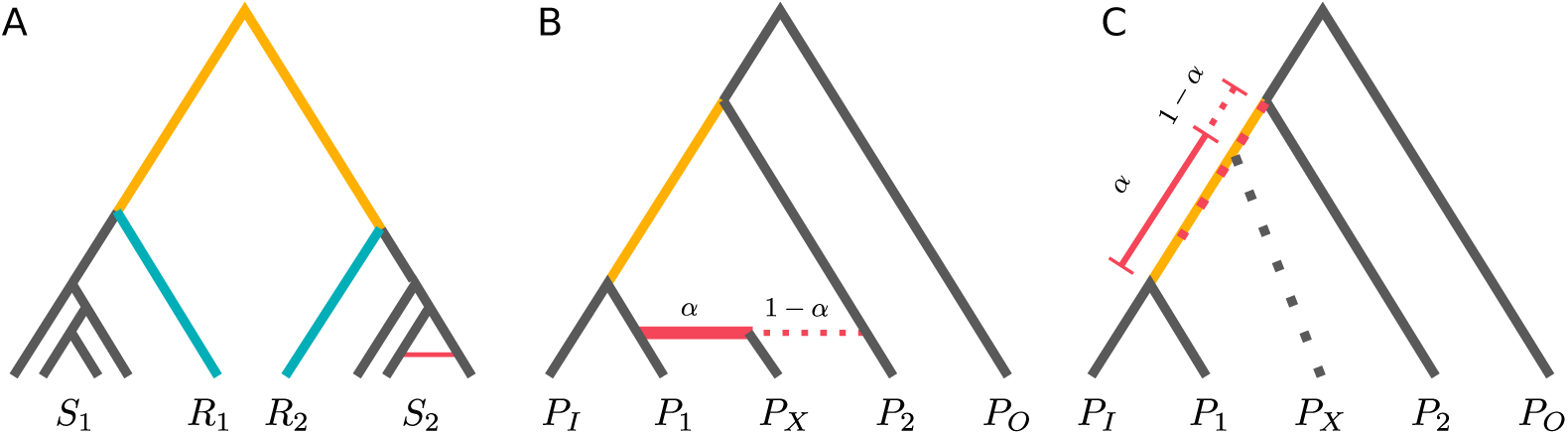
Applications of *F*_4_: A: Visualization of rank test to estimate the number of admixture events. *F*_4_(*S*_1_, *R*_1_, *S*_2_, *R*_2_) measures a branch absent from the phylogeny and should be zero for all populations from *S*_1_ and *S*_2_. B: Model underlying admixture ratio estimate [10]. *P_X_* splits up, and the mean coalescence time of *P_X_* with *P_I_* gives the admixture proportion. C: If the model is violated, *α_X_* measures where on the internal branch in the underlying genealogy *P_X_* (on average) merges

#### Admixture proportion

The second application is by comparing branches between closely related populations to obtain an estimate of mixture proportion, or how much two focal populations correspond to an admixed population. [10]:

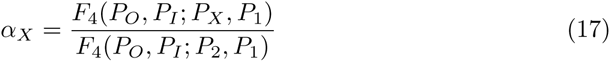

Here, *P_X_*, is the population whose admixture proportion we are estimating, *P*_1_ and *P*_2_ are the potential contributors, where we assume that they contribute with proportions *α_X_* and 1 − *α_X_*, respectively. and *P_O_*, *P_I_* are reference populations with no direct contribution to *P_X_* (see Figure 5B). *P_I_* has to be more closely related to one of *P*_1_ or *P*_2_ than the other, and *P_O_* is an outgroup.

The canonical way [5] to interpret this ratio is as follows: the denominator is the branch length from the common ancestor population from *P_I_* and *P*_1_ to the common ancestor of *P_I_* with *P*_2_. (Figure 5C, yellow line), The numerator has a similar interpretation as an internal branch (red dotted line). In an admixture scenarios, (Figure 5B, this is not unique, and is replaced by a linear combination of lineages merging at the common ancestor of *P_I_* and *P*_1_ (with probability *α_X_*), and lineages merging at the common ancestor of *P_I_* with P_2_ (with probability 1 − *α_X_*).

Thus, a more general interpretation is that *α_X_* measures how much closer the common ancestor of *P_X_* and *P_I_* is to the common ancestor of *P_I_* and *P*_1_ and the common ancestor of *P_I_* and *P*_2_, indicated by the gray dotted line in Figure 5B. This quantity is defined also when the assumptions underlying the admixture test are violated, and if the assumptions are not carefully checked, might lead to misinterpretations of the data. In particular, *α_X_* is well-defined in cases where no admixture occurred, or in cases where either of *P*_1_ and *P*_2_ did not experience any admixture.

Furthermore, it is evident from Figure 5 that if all populations are sampled at the same time, 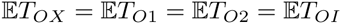, and therefore,

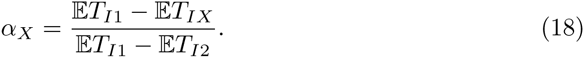

Thus,

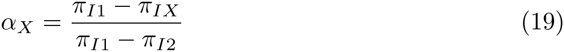

is another estimator for *α_X_* that can be used even if no outgroup is available. We compared Equations 17 and 19 for varying admixture proportions in Figure 4C using the mean absolute error in the admixture proportion. Both estimators perform very well, but we find that (19) performs slightly better in cases where the admixture proportion is low. However, in most cases this minor improvement possibly does not negate the drawback that Equation 19 is only applicable when populations are sampled at the same time.

## Structure models

For practical purposes, it is useful to know how the admixture tests perform under demographic models different from population phylogenies and admixture graphs, and in which cases the assumptions made for the tests are problematic. In other words, under which demographic models is population structure well-approximated by a tree? Equation 5 allows us to derive expectations for *F*_3_ and *F*_4_ under a wide variety of models of population structure (Figure 6). The simplest case is that of a single panmictic population. In that case, all *F*-statistics have an expectation of zero, consistent with the assumption that no structure and therefore no population phylogeny exists. Under island models, *F*_4_ is also zero, and *F*_3_ is inversely proportional to the migration rate. Results are similar under a hierarchical island model, except that the number of demes has a small effect. This corresponds to a population phylogeny that is star-like and has no internal branches, which is explained by the strong symmetry of the island model. Thus, looking at different *F*_3_ and *F*_4_-statistics may be a simple heuristic to see if data is broadly consistent with an island model; if *F*_3_-values vary a lot between populations, or if *F*_4_ is substantially different from zero, an island model might be a poor choice. When looking at a finite stepping stone model, we find that *F*_3_ and *F*_4_ are both non-zero, highlighting that *F*_4_ (and the ABBA-BABA-*D*-statistic) is susceptible to migration between any pair of populations. Thus, for applications, *F*_4_ should only be used if there is good evidence that gene flow between some pairs of the populations was severely restricted. A hierarchical stepping stone model, where demes are combined into populations, is the only case besides the admixture graph where *F*_3_ can be negative. This effect indicates that admixture and population structure models may be the two sides of the same coin: we can think of admixture as a (temporary) reduction in gene flow between individuals from the same population. Finally, for a simple serial founder model without migration, we find that *F*_3_ measures the time between subsequent founder events.

**Figure 6.**
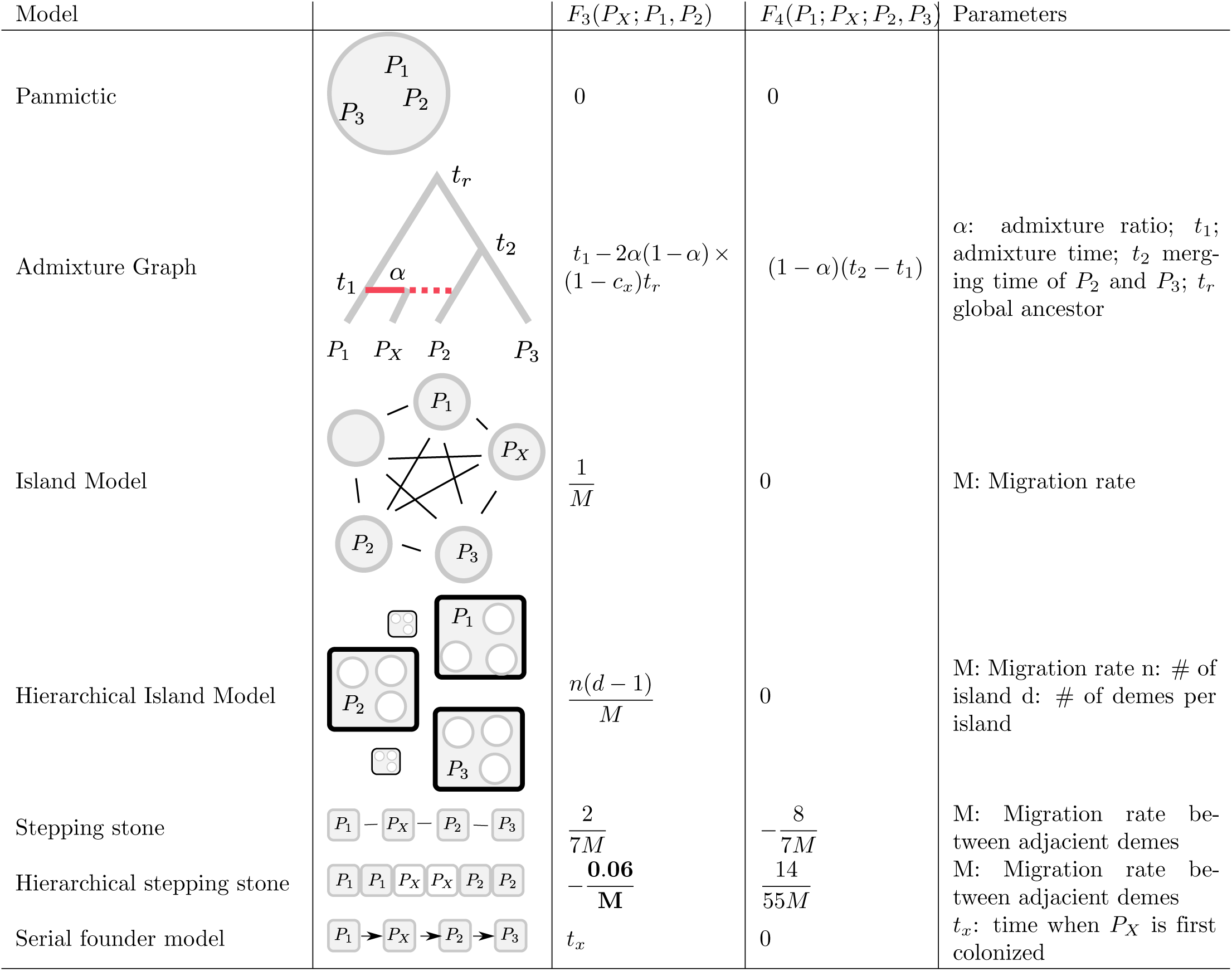
Expectations for *F*_3_ and *F*_4_ under select models.

## Conclusions

We showed that there are three main ways to interpret *F*-statistics: First, we can think of them as the branches in a population phylogeny. Second, we can think of them as the shared drift, or paths in an admixture graph. And third, we can think of them in terms of coalescence times and the lengths of the internal branches of gene genealogies. This last interpretation allows us to make the connection to the ABBA-BABA-statistic explicit, and allows us to investigate the behavior of the *F*-statistics under arbitrary demographic models.

If we have indices for two, three and four populations, should there be corresponding quantities for five or more populations(e.g. [36])? Two of the interpretations speak against this possibility: First, a population phylogeny can be fully characterized by internal and external branches, and it is not clear how a five-population statistic could be written as a meaningful branch length. Second, we can write all *F*-statistics in terms of four-individual trees, but this is not possible for five samples. This seems to suggest that there may not exist a five-population statistic as general as the *F*-statistics we discussed here, but they will still be valid for questions pertaining to a very specific demographic model [36].

A well-known drawback of *F*_3_ is that it may have a positive expectation under some admixture scenarios [5]. Here, we showed that *F*_3_ is positive if and only if the branch supporting the population tree is longer than the two branches discordant with the population tree. Note that this is (possibly) distinct from the probabilities of tree topologies, although the average branch length of the internal branch in a topology, and the probability of that topology may frequently very correlated. Thus, negative *F*_3_-values indicate that individuals from the admixed population are more likely to coalesce with individuals from the two other populations, than with other individuals from the same population!

Overall, when *F*_3_ is applicable, it is remarkably robust to population structure, requiring rather strong substructure to yield false-positives. Thus, it is a very striking finding that in many applications to humans, negative *F*_3_-values are commonly found [4,5], indicating that for most human populations, the majority of markers support a discordant gene tree, which suggests that population structure and admixture are widespread and that population phylogenies are poorly suited to describe human evolution.

Ancient population structure was proposed as possible confounder for the *D* and *F*_4_-statistics [10]. Here, we show that non-symmetric population structure such as in stepping stone models can lead to non-zero *F*_4_-values, showing that both ancestral and persisting population structure may result in false-positives when the statistics are applied in an incorrect setting.

Furthermore, we showed that the *F*-statistics can be seen as a special case of a tree-metric, and that both *F*_3_ and *F*_4_ can be interpreted, for arbitrary tree metrics, as tests for properties of phylogenetic trees.

From this perspective, it is worth re-raising the issue pointed out by Felsenstein [21], how and when allele-frequency data should be transformed for within-species phylogenetic inference. While *F*_2_ has become a *de facto* standard, which, as we have shown, leads to useful interpretations, the *F*_3_ and *F*_4_-tests can be used for arbitrary tree metrics, and different transformations of allele frequencies might be useful in some cases.

But it is clear that, when we are applying *F*-statistics, we are implicitly using phylogenetic theory to test hypotheses about simple phylogenetic networks [37].

This close relationship provides ample opportunities for interaction between these currently diverged fields: Theory [37, 38] and algorithms for finding phylogenetic networks such as Neighbor-Net [39] may provide a useful alternative to tools specifically developed for allele frequencies and *F*-statistics [5-7], particularly in complex cases. On the other hand, the tests and different interpretations described here may be useful to test for treeness in other phylogenetic applications, and the complex history of humans may provide motivation to further develop the theory of phylogenetic networks, and stress its usefulness for within-species demographic analyses.

## Acknowledgements

I would like to thank Heejung Shim, Choongwon Jeong, Evan Koch, Lauren Blake, Joel Smith and John Novembre for helpful comments and discussions.

## Methods

### Equivalence of drift interpretations

First, we show that *F*_2_ can be interpreted as the difference in variance of allele frequencies (Figure 1C):

As in the Results section, let *P_i_* denote a population with allele frequency, sample size and sampling time with *p_i_*, *n_i_* and *t_i_*, respectively. Then, for *t*_o_ < *t_t_*:

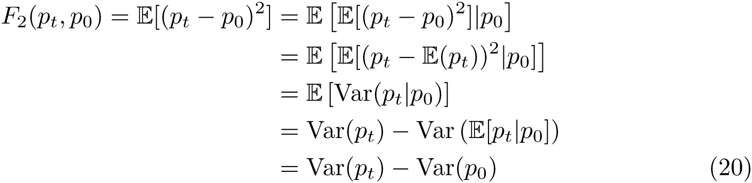

Here, we used 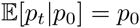 on lines two and five (which holds if there is no mutation, no selection and *P_t_* is a descendant of *P*_0_). The fourth line is obtained using the law of total variance. It is worth noting that this result holds for any model of genetic drift where the expected allele frequency is the current allele frequency (the process describing the allele frequency is a martingale). For example, this this interpretation of *F*_2_ holds also if we model genetic drift as a Brownian motion.

#### A heterozygosity model

The interpretation of *F*_2_ in terms of the decay in heterozygosity and identity by descend can be derived elegantly using duality between the diffusion process and the coalescent: Let again *t*_0_ < *t_t_* Furthermore, let *f* be the probability that two individuals sampled at time *t_t_* have coalesced at time *t*_0_.

Then,

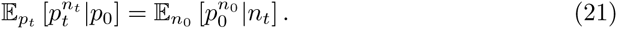

This equation is due to Tavaré [40], who also provided the following intuition: Given we sample *n_t_* individuals at time *t_t_* let *E* denote the event that all individuals carry allele *x*, conditional on allele *x* having frequency *p*_0_ at time *t*_0_. There are two components to this: First, the frequency will change between *t*_0_ and *t_t_*, and then we need all *n_t_* sampled individuals to carry *x*.

In a diffusion framework, we can write

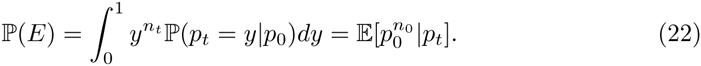

On the other hand, we may argue using the coalescent: For *E* to occur, all *n*_1_ samples need to carry the *x* allele. At time *t*_0_, they had *n*_0_ ancestral lineages, who all carry *x* with probability *p*_0_. Therefore,

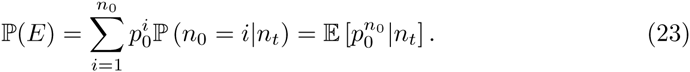

Equating (22) and (23) yields Equation 21.

In the present case, we are most interested in the cases of *n_t_* = 1, 2, since:

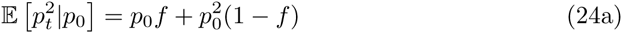

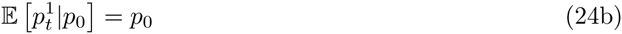

To derive an expression for *F*_2_, we start by conditioning on the allele frequency *p*_0_,

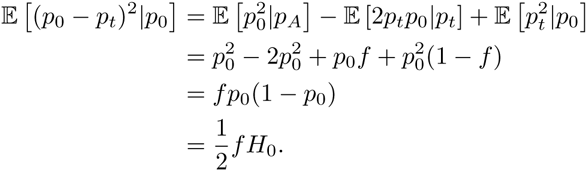

Where *H*_0_ = 2*p*_0_(1 – *p*_0_) is the heterozygosity. Integrating over *p*_0_ yields:

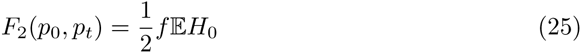

and we see that *F*_2_ increases as a function of *f* (Figure 1E). This equation can also be interpreted in terms of probabilities of identity by descent: *f* is the probability that two individuals are identical by descent in *P_t_* given their ancestors were not identical by descent in *P*_0_, and 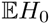 is the probability two individuals are not identical by descent in *P*_0_. Thus, *F*_2_ is half the probability of the event that two individuals in *P_t_* are identical by descent, and they were not in *P*_0_.

Furthermore, 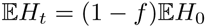 (Equation 3.4 in [28]) and therefore

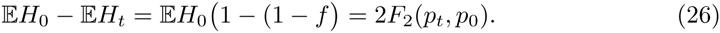

which shows that *F*_2_ measures the decay of heterozygosity (Figure 1C). A similar argument was used by in [7] to estimate ancestral heterozygosities using *F*_2_ and to linearize *F*_2_.

#### Two populations

*F*_2_ in terms of the difference in expected and observed heterozygosity follows directly from the result from [19], which was obtained by considering the genotypes of all possible matings in the two subpopulations, and the variance case follows directly because 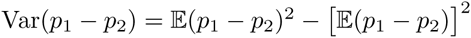, but 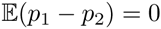. Lastly, we relate *F*_2_ to *F_ST_* by using the definition of *F*_2_ as a variance in the definition of *F_ST_*: 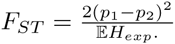

#### Covariance interpretation

To see how *F*_2_ can be interpreted as a covariance between two individuals from the same population, define *X_i_* and *X_j_* as indicator variables that two individuals from the same population sample have the A allele, which has frequency *p*_1_ in one, and and *p*_2_ in the other population. If we are equally likely to pick from either population,

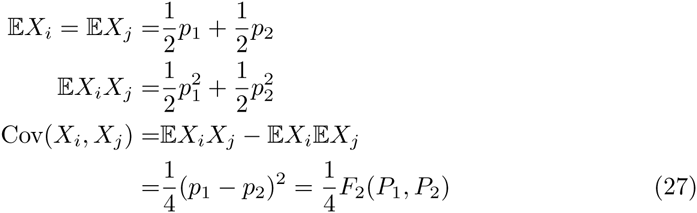

The expectations can be interpreted that we pick a population, and then with probability equal to the allele frequency an individual will have the *A* allele. The joint expectation is similar, except we need two individuals.

### Derivation of *F*_2_ for gene trees

To derive equation (5), we start by considering *F*_2_ for two samples of size one, express *F*_2_ for arbitrary sample sizes in terms of individual-level *F*_2_, and obtain a sample-size independent expression by letting the sample size *n* go to infinity.

In this framework, we assume that mutation is rare such that there is at most one mutation at any locus. In a sample of size two, let the genotypes of the two haploid individuals be denoted as *I*_1_, *I*_2_. *I_i_* ∈ {0, 1} and *F*_2_(*I*_1_, *I*_2_) = 1 implies *I*_1_ = *I*_2_, whereas *F*_2_(*I*_1_, *I*_2_) = 0 implies *I*_1_ ≠ *I*_2_. We can think of *F*_2_(*I*_1_, *I*_2_) as an indicator random variable with parameter equal to the branch length between *I*_1_ and *I*_2_, times the probability that a mutation occurs on that branch.

Now, replace *I*_1_ with a sample 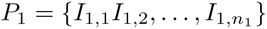. The sample allele frequency is 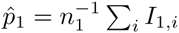. And the sample-*F*_2_ is

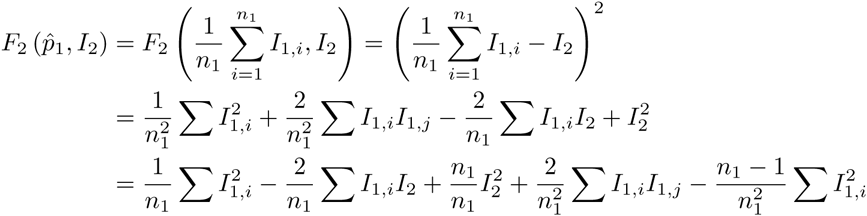

The first three terms can be grouped into *n*_1_ terms of the form *F*_2_(*I*_1_,*_i_*, *I*_2_), and the last two terms can be grouped into 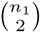 terms of the form *F*_2_(*I*_1,_*_i_ I*_1_,*_j_*), one for each possible pair of samples in *P*_1_.

Therefore,

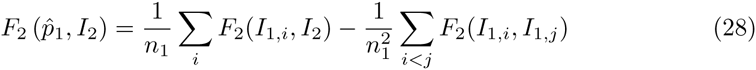

where the second sum is over all pairs in *P*_1_.

As 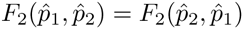, we can switch the labels, and obtain the same expression for population *P*_2_ = {*I*_2_,*_i_*, *i* = 0,…, *n*_2_} Taking the average over all *I*_2_,*_j_* yields

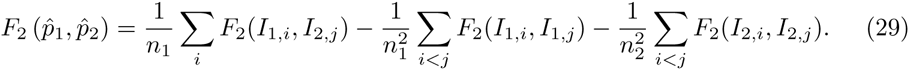

Thus, we can write *F*_2_ between the two populations as the average number of differences we see between individuals from different populations, minus some terms including differences *within* each sample.

Equation 29 is quite general, making no assumptions on where samples are placed on a tree. In a coalescence framework, it is useful to make the assumptions that all individuals from the same population have the same branch length distribution, i.e. 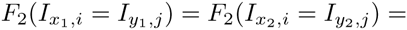 for all pairs of samples (*x*_1_, *x*_2_) and (*y*_1_, *y*_2_) from populations *P_i_* and *P_j_*. Secondly, we assume that all samples correspond to the leaves of the tree, so that we can estimate branch lengths in terms of the time to a common ancestor *T_ij_*. Finally, we assume that mutations occur at a constant rate of *θ*/2 on each branch. Taken together, these assumptions imply that 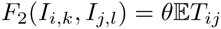 for all individuals from populations *P_i_*, *P_j_*, this simplifies to

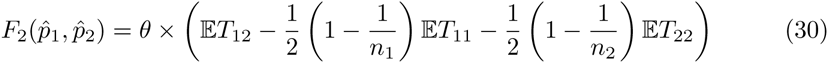

which, for the cases of *n* =1, 2 was also derived by Petkova [41]. In most applications, we wish to calculate *F*_2_ per segregating site in a large sample. As the expected number of segregating sites is 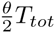, we can follow [24,41] and take the limit where *θ* → 0:

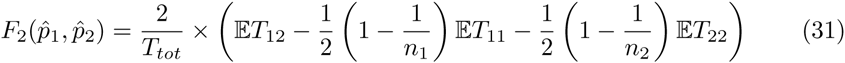

to obtain an expression independent of the mutation rate. In either of these equations, we can see 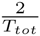 or *θ* as a constant of proportionality that is the same for all statistics calculated from the same data. Since we are either interested in the relative magnitude of *F*_2_, or whether a sum of *F*_2_-values is different from zero, this constant has no impact on inference.

Furthermore, we can obtain a population-level statistic by taking the limit when the number of individuals per sample *n*_1_ and *n*_2_ go to infinity:

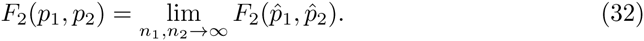

This yields Equation 5. Using *θ* as the constant of proportionality, we find that

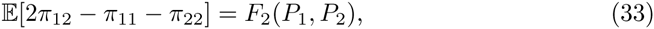

leading to the estimator given in 6.

It is straightforward to check that this estimator is equivalent to that given by Reich *et al.* [4]:

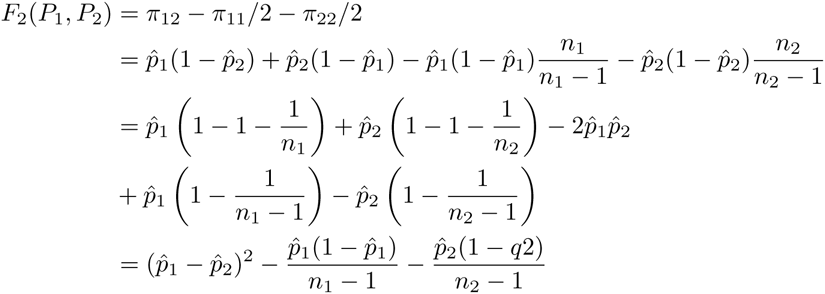

which is Equation 10 in the Appendix of [4].

### Four-point-condition and *F*_4_

We prove the statement that for any tree, two of the three possible *F*_4_ values will be equal, and the last will be zero. First, notice that permuting one of the two pairs only changes the sign of the statistic, i.e.

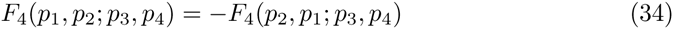

Using *F*_2_ as a tree-metric, the four-point condition [17] can be written as

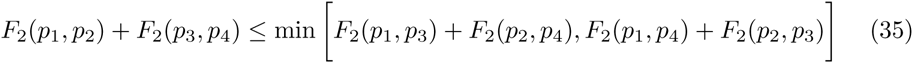

which holds for any permutations of the samples.

Applying this to the first two and last two terms on the right-hand-side in equation 14b yields

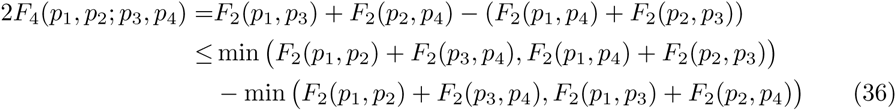

The four-point condition states that two of the sums of disjoint *F*_2_ statistics need to be identical, and the third one should be less or equal than that. This gives us four cases to evaluate (36) under:

1. If *F*_2_(*p*_1_, *p*_2_) + *F*_2_(*p*_3_, *p*_4_) is smallest: (36) is zero
2. If *F*_2_(*p*_1_, *p*_3_) + *F*_2_(*p*_2_, *p*_4_) is smallest: (36) is *F*_4_(*p*_1_, *p*_4_; *p*_2_, *p*_3_) > 0
3. If *F*_2_(*p*_1_, *p*_4_) + *F*_2_(*p*_2_, *p*_3_) is smallest: (36) is −*F*_4_(*p*_1_, *p*_4_; *p*_2_, *p*_3_) < 0
4. All sums of *F*_2_ are equal: (36) is zero

If the *F*_2_ are not all equal, then for each *F*_4_ with distinct pairs, one of conditions 2-4 is true, and we see that indeed one will be zero and the other two will have the same absolute value.

### Derivation of *F* under select models

Here, we use Equation 5 together with Equations 10b and 14b to derive expectations for *F*_3_ and *F*_4_ under some simple models.

### Panmixia

Under panmixia with arbitrary population size changes, *P*_1_ and *P*_2_ are taken from the same pool of individuals and therefore *T*_12_ = *T*_11_ = *T*_22_, 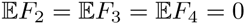.

#### Island models

A (finite) island model has *D* subpopulations of size 1 each. Migration occurs at rate *M* between subpopulations. It can be shown [42] that 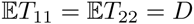. 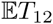 satisfies the recursion

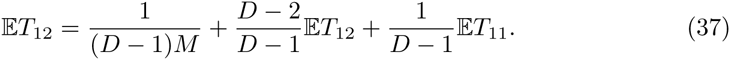

with solution 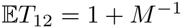. This results in the equation in figure 6. The derivations for the hierarchical island models is marginally more complicated, but similar. It is given in [43].

#### Admixture models

These are the model for which the *F*-statistics were originally developed. Many details, applications, and the origin of the path representation are found in [5]. For simplicity, we look at the simplest possible tree of size four, where *P_X_* is admixed from *P*_1_ and *P*_2_ with contributions *α* and *β* = (1 − *α*), respectively. We assume that all populations have the same size, and that this size is one. Then,

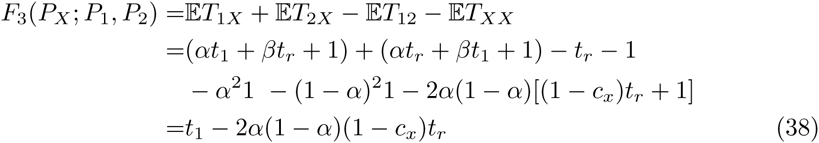

Here, *c_x_* is the probability that the two lineages from *P_X_* coalesce before the admixture event.

Thus, we find that *F*_3_ is negative if

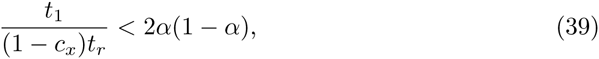

which is more likely if *α* is large, the admixture is recent and the overall coalescent is far in the past.

For *F*_4_, we have, omitting the within-population coalescence time of 1:

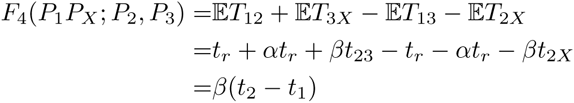

#### Stepping-stone models

For the stepping stone models, we have to solve the recursions of the Markov chains describing the location of all lineages in a sample of size 2. For the standard stepping stone model, we assumed there were four demes, all of which exchange migrants at rate *M*. This results in a Markov Chain with the following five states: i) lineages in same deme ii) lineages in demes 1 and 2, iii) lineages in demes 1 and 3, lineages in demes 1 and 4 and v) lineages in demes 2 and 3. Note that the symmetry of this system allows us to collapse some states. The transition matrix for this system is

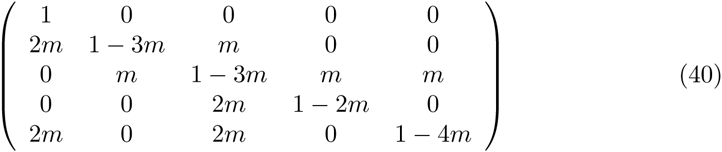

We can end the system once lineages are in the same deme, as the time to coalescence time is independent of the deme in isotropic migration models [42], and cancels from the *F*-statistics.

Therefore, we can find the vector *v* of the expected time until two lineages using standard Markov Chain theory by solving *v* = (**I** − **T**)^−1^)**1**, where **T** is the transition matrix involving only the transitive states in the Markov chain (all but the first state), and **1** is a vector of ones.

Finding the expected coalescent time involves solving a system of 5 equations. The terms involved in calculating the *F*-statistics (Table 1) are the entries in *v* corresponding to these states.

The hierarchical case is similar, except there are 6 demes and 10 equations. Representing states as lineages being in demes (same), (1,2), (1,3), (1,4), (1,5), (1,6), (2,3), (2,4), (2,5), (3,4).

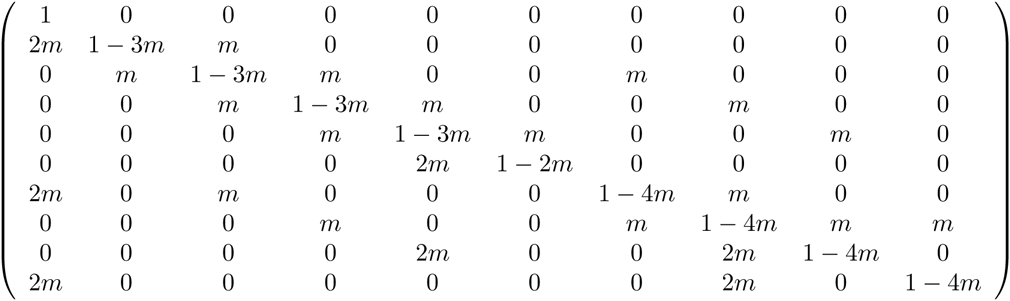

And we can solve the same equation as in the non-hierarchical case to get all pairwise coalescence times. Then, all we have to do is average the coalescence times over all possibilities. E.g.

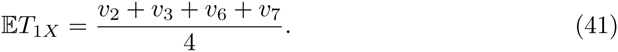

For *F*_4_, we assume that demes 1 and 2 are in *P*_1_, demes 3 and 4 in *P_X_* and demes 5 and 6 correspond to *P*_2_ and *P*_3_, respectively.

We average the two left demes to *P*_1_, the two right demes to *P*_2_ and the two middle demes to *P_X_*. The

#### Range expansion model

We use a range expansion model with no migration [44]. Under that model, we assume that samples *P*_1_ and *P*_2_ are taken from demes *D*_1_ and *D*_2_, with *D*_1_ closer to the origin of the expansion, and populations with high ids even further away from the expansion origin. Then 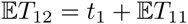, where 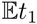 is the time required for a lineage sampled further away in the expansion to end up in *D*_1_. (Note that *t*_1_ only depends on the deme that is closer to the origin). Thus, for three demes,

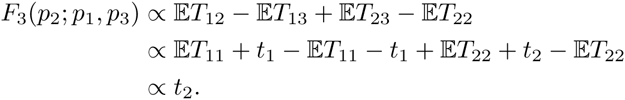

and

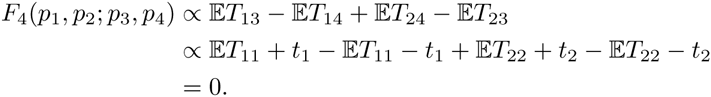

More interesting is

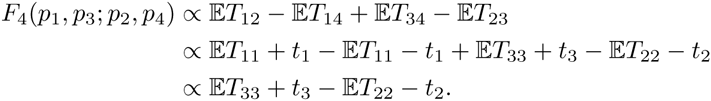

## Simulations

Simulations were performed using ms [45]. Specific commands used are

~~~
ms 1201 100 -t 10 -I 13 100 100 100 100 100 100 100 100 100 100 100 100 1 -ej 0.01 2 1 -ej 0.02 3 1 -ej 0.04 4 1 -ej 0.06 5 1 -ej 0.08 6 1 -ej 0.10 7 1 -ej 0.12 8 1 -ej 0.14 9 1 -ej 0.16 10 1 -ej 0.16 11 1 -ej 0.3 12 1 -ej 0.31 13 1
~~~

for the outgroup-*F*_3_-statistic (Figure 4A),

~~~
ms 301 100 -t 10 -I 4 100 100 100 1 -es 0.001 2 $ALPHA -ej 0.03 2 1 -ej 0.03 5 3 -ej 0.3 3 1 -ej 0.31 4 1
~~~

for Figure 4B, where the admixture proportion `$ALPHA` was varied in increments of 0.025 from 0 to 0.5, with 200 data sets generated per `$ALPHA`.

Lastly, data for Figure 4C was simulated using

~~~
ms 501 100 -t 50 -r 50 10000 -I 6 100 100 100 100 100 1 -es 0.001 3 $ALPHA -ej 0.03 3 2 -ej 0.03 7 4 -ej 0.1 2 1 -ej 0.2 4 1 -ej 0.3 5 1 –ej 0.31 6 1
~~~

Here, the admixture proportion `$ALPHA` was varied in increments of 0.1 from 0 to 1, again with 200 data sets generated per `$ALPHA`.

*F*_3_ and *F*_4_-statistics were calculated using the implementation from [6].

---

**Figure S1.**
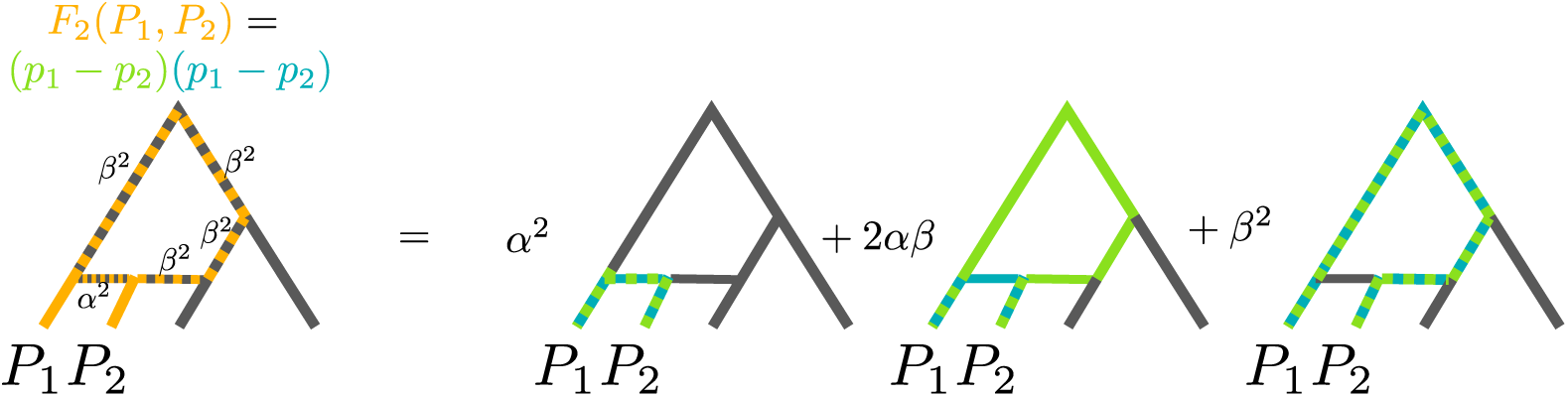
Path interpretation of *F*_2_: *F*_2_ is interpreted as two possible paths from *P*_1_ to *P*_2_, which we color green and blue, respectively. With probability *α*, a path takes the left admixture edge, and with probability *β* = 1 − *α*, the right one. The dotted lines give the overlap of the two paths, conditional on which admixture edge they take, and the result is summarized as the weighted sum of branches in the left-most tree. For a more detailed explanation, see [5].

**Figure S2.**
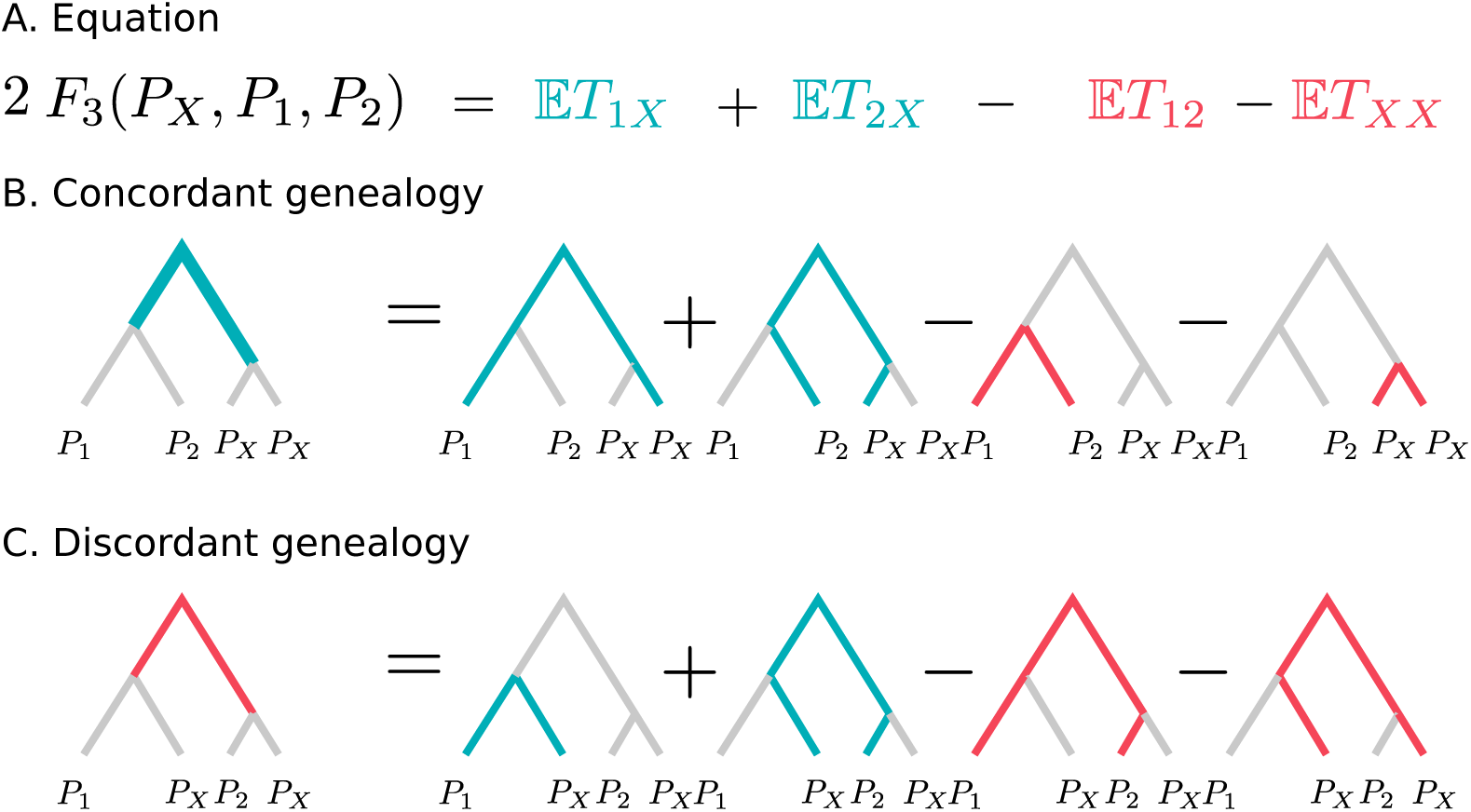
Schematic explanation how *F*_3_ behaves conditioned on gene tree. Blue terms and branches correspond to positive contributions, whereas red branches and terms are subtracted. Labels represent individuals randomly sampled from that population. We see that external branches cancel out, so only the internal branches have non-zero contribution to *F*_3_. In the concordant genealogy (Panel B), the contribution is positive (with weight 2), and in the discordant genealogy (Panel C), it is negative (with weight 1). The mutation rate as constant of proportionality is omitted.

**Figure S3.**
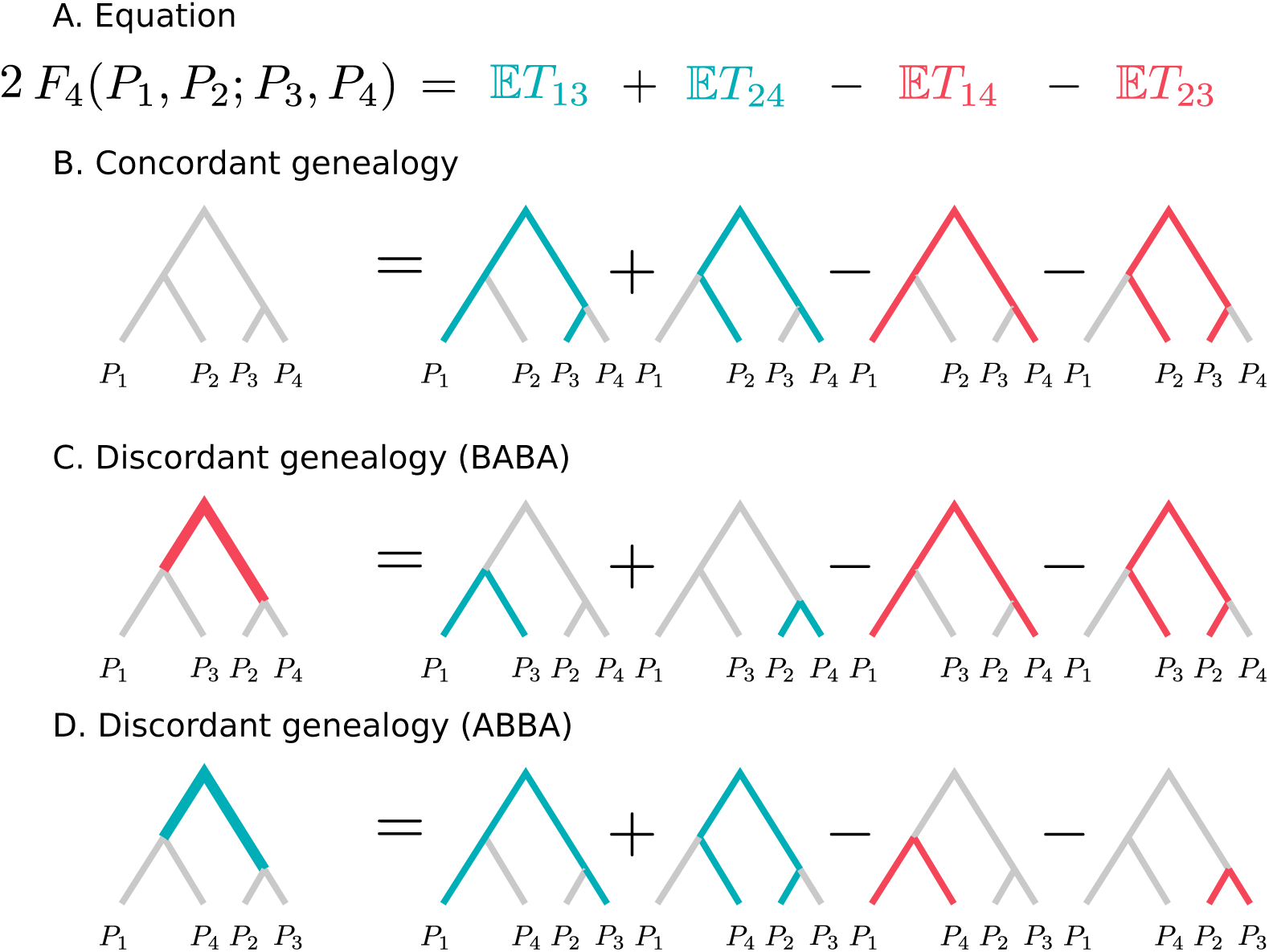
Schematic explanation how *F*_4_ behaves conditioned on gene tree. Blue terms and branches correspond to positive contributions, whereas red branches and terms are subtracted. Labels represent individuals randomly sampled from that population. We see that all branches cancel out in the concordant genealogy (Panel B), and that the two discordant genealogies contribute with weight +2 and −2, respectively The mutation rate as constant of proportionality is omitted.

